# Benchmarking non-targeted metabolomics using yeast derived libraries

**DOI:** 10.1101/2020.10.06.319160

**Authors:** Evelyn Rampler, Gerrit Hermann, Gerlinde Grabmann, Yasin El Abiead, Harald Schoeny, Christoph Baumgartinger, Thomas Köcher, Gunda Koellensperger

## Abstract

Non-targeted analysis by high-resolution mass spectrometry (HRMS) is the essential discovery tool in metabolomics. Up to date, standardization and validation remain a challenge. Community wide accepted, cost-effective benchmark materials are lacking. In this work, we propose yeast (*Pichia pastoris*) extracts, derived from fully controlled fermentations for this purpose. We established an open-source metabolite library of > 200 metabolites, reproducibly recovered in ethanolic extracts by orthogonal LCHRMS methods, different fermentations (over three years) and different laboratories. More specifically, compound identification was based on accurate mass, matching retention times, and MS/MS spectra as compared to authentic standards and internal databases. The library includes metabolites from the classes of 1) organic acids and derivatives (2) nucleosides, nucleotides and analogues, (3) lipids and lipid-like molecules, (4) organic oxygen compounds, (5) organoheterocyclic compounds, (6) organic nitrogen compounds and (7) benzoids at expected concentrations ranges of sub-nM to µM. As yeast is a eukaryotic organism, key regulatory elements are highly conserved between yeast and all annotated metabolites were also reported in the Human metabolome data base (HMDB). A large fraction of metabolites was found to be stable for several years when stored at −80°C. Thus, the yeast benchmark material enabled not only to test for the chemical space and coverage upon method implementation and developments, but enabled in-house routines for instrumental performance tests. Finally, the benchmark material opens new avenues for batch to batch corrections in large scale non-targeted metabolomics studies.

## Introduction

Non-targeted analysis (NTA) by high resolution mass spectrometry (HRMS) is a prime example of an innovative measurement practice and the key to discoveries in many applications such as different ‘omics’ areas^1^, environmental protection^2^, and food safety^3^. NTA will not replace targeted analysis, but emerged as a complementing cost-effective discovery tool. Especially omics-scale research profits from the improved coverage and the lower costs of analysis provided by HRMS. The common challenge in all fields of application is to create reliability of data and to find ways of harmonization especially in large scale multicenter studies^4–6^. As a matter of fact, validation practices and guidelines from targeted analysis cannot be simply applied. Thus, validation in NTA is significantly less developed and defined. This fact, together with the wide acceptance of the NTA tool set calls for new strategies of standardization, quality control measures, and metrics for evaluation. Joint efforts towards harmonized NTA protocols and definition of a minimum of quality requirements are of paramount importance in this endeavor.

Metabolomics is one of the key applications of NTA. The metabolomics standardization initiative (MSI) of the metabolomics society^7^ has worked intensively on definitions and guidelines considering all steps of the NTA analytical process since many years. This includes defining the analytical task, sampling/analysis of data standards, data evaluation, and reporting^8,9^. NTA was addressed with regard to the annotation of metabolites and their relative quantification at all levels of analysis. MSI recommends that all researchers define the level of identification, a common name, and a structural code (e.g., InChI or SMILES) in their publications^10^. MSI also highlights the need to submit the data to open-access repositories such as MetaboLights^11^ to provide clarity of NTA data. The impact of this initiative on all levels of NTA was massive and significantly improved the harmonization in the field^5^.

Reference materials meeting all metrological stringent criteria of full traceability are scarce in metabolomics and in life sciences in general. One reason might be the rather poor acceptance of reference materials as heir extensive use in large scale omics-type of measurement campaigns increase costs singnificantlys. In fact, the list of available reference materials for metabolomics is very short. In large scale metabolomics studies, the concept of a pooled sample for quality control has gained worldwide acceptance allowing to correct for intra- and inter-batch variations and to accomplish MS/MS measurements required for annotation^5^. However, in multicenter studies as envisioned in clinical metabolomics, it is not straightforward to produce pooled samples in sufficient amounts. Only very recently (June 2020), the first untargeted metabolomics study on a large scale basis comparing three pooled human plasma reference materials (Qstd3, 211 CHEAR, NIST1950) was reported^12^.

In other omics fields such as proteomics, the concept of cheap and easily accessible benchmark materials prevailed. HeLa cell extracts have become the gold standard for benchmarking instrument performance and proof-of-principle experiments upon introduction of new analytical methods^13–17^.

In proteomics, many laboratories and instrument manufacturers resort to tryptic digests of protein extracts from HeLa cells to check performance^16^, proof a method fit-for-purpose, benchmark new proteomics workflows^13^ or show significant technological progress^18^. All metrics are established for quality control, protein identification numbers, and reported compound areas.

Currently, there is no such commonly accepted low cost biological matrix material in metabolomics. In this work, we explore yeast as a potential benchmark material for metabolomics. Yeasts are industrially important cell factories easy to cultivate in short time and large populations using inexpensive media. Despite the phylogenetical distance, a number of key regulatory elements are highly conserved between yeast and humans^19^. In the field of metabolomics, ^13^C enriched yeast has become widely accepted as resource for ^13^C internal standards. Controlled growth conditions of *Pichia pastoris* are ideal to produce ^13^C labeled metabolites with efficiencies higher than 99% leading the simultaneous production of hundreds of biological relevant labeled metabolites^20–22^. These highly enriched compounds enabled absolute quantification for a wide range of metabolites and lipids^20,23–25^ as well as validation of new software tools for non-targeted data evaluation such as the METLIN platform (using fragment matches of labeled and non-labeled metabolite pairs)^26^.

In this work, we propose the use of ethanolic yeast extracts from *Pichia pastoris in vivo* fermentation as stable and cheap controls for non-targeted metabolomics workflows. Evidently, a benchmark material cannot replace certified reference materials, but as in proteomics, it could play a significant role facilitating method development and validation. A first crucial step in this direction is our reported metabolite library.

## Material and Methods

### Standards and solvents

Metabolite standards were purchased from Sigma Aldrich (Vienna, Austria) or Carbosynth (Berkshire, UK). Lipid Standards were obtained from Avanti Polar Lipids, Inc. (Alabaster, Alabama, USA), Sigma Aldrich (Vienna, Austria) or Carbosynth (Berkshire, UK). All lipid and metabolite standards were weighed, dissolved in an appropriate solvent (mostly methanol or water) and a multi-metabolite as well as a multi-lipid mix were prepared. The standard reference material (SRM) 1950 Metabolites in Frozen Human Plasma was purchased from the National Institute of Standards & Technology (NIST) (Gaithersburg, USA). Acetonitrile (ACN), isopropanol (IPA), methanol (MeOH), and water were of LC-MS grade and ordered from Fisher Scientific (Vienna, Austria) or Sigma Aldrich (Vienna, Austria). Ammonium bicarbonate, ammonium formate, and ammonium hydroxide were ordered as LC-MS grade eluent additives from Sigma Aldrich. Formic acid was also of LC-MS grade and obtained from VWR International (Vienna, Austria).

### Production of ethanolic yeast extracts

*Pichia pastoris* (Guillierm.) Phaff 1956 (*Komagataella phaffii Kurtzman*)^27^ was cultivated in a New Brunswick BioFlo 310 fed-batch fermenter for 38 h (Eppendorf, Hamburg, Germany) with full control over the input variables in terms of glucose as carbon source (Cambridge Isotope laboratories), pH, temperature, and oxygenation. Process monitoring was facilitated by online measurement of pH, temperature, and dissolved oxygen. Offline assessment of optical density at 600 nm (OD600) and optical cell counting was performed at several time points. The cells were fermented until an OD600 of 12. At the end of the process, the biomass was quenched in 60% methanol (v/v) at −30 °C and subsequently extracted in boiling 80% ethanol (v/v) for metabolites. Finally, the ethanolic extract was aliquoted and dried in a vacuum centrifuge (Genevac EZ2). Aliquots of the *Pichia pastoris* ethanolic extracts, derived from ∼2 billion yeast cells (corresponding to 20 mg of cell dry weight), were dried in 15 mL Falcon tubes and stored at −80 °C prior to analysis.

### LC-MS analysis and data analysis of yeast extracts

#### Untargeted Metabolomics

Samples of dried ethanolic yeast extract (derived from two different batches: 11/2017, 05/2018) were dissolved in 2.5 mL MiliQ water (Advantage Q10, Merck) and sample batch 05/2019 was filled up to a volume of 5 mL with MiliQ water. An aliquot of 100 µL was transferred into an Eppendorf tube and centrifuged at 4 °C for 10 min at 15 000 RCF. 50 µL of the supernatant was diluted 1:1 (v/v) with acetonitrile or 0.1% formic acid, for HILIC or RPLC measurements, respectively. 10 μL of each sample were pooled and used as a quality control (QC) sample. Samples were randomly assigned into the autosampler and metabolites were separated on a SeQuant ZIC-pHILIC HPLC column (Merck, 100 x 2.1 mm; 5 µm with guard column) or a RP-column (Waters, ACQUITY UPLC HSS T3 150 x 2.1 mm; 1.8 μm with VanGuard column) with a flow rate of 100 µL min^−1^ delivered through an Ultimate 3000 HPLC system (Thermo Fisher Scientific, Germany). The stepwise gradient for HILIC analysis (adapted from Wernisch and Pennathur^28^) involved starting conditions of 90% A (100% ACN), ramp to 25% B (25 mM ammonium hydrogen carbonate, pH 8) within 6 min, 2 min hold at 25% B, from 8 to 21 min a ramp to 60% B was applied, switching to 80% B at 21.5 min followed by a flushing (21.5-26 min: 80% B) and reequilibration step (26.1-35 at 10% B). The gradient for RPLC analysis involved a linear ramp up time of 20 min starting with 99% A (0.1% formic acid) to 60% B (0.1% formic acid in ACN) followed by 5 min hold (21-26 min) at 90% B and a reequlibration step (26.1-36 min: 1% B). All samples were analyzed by both HILIC and RP separation followed by ESI-HRMS in polarity switching mode in the mass range 60-900 m/z. Sample spectra were acquired by data-dependent high-resolution tandem mass spectrometry on a Q Exactive Focus (Thermo Fisher Scientific, Germany). Ionization potential was set to +3.5/-3.0 kV, the sheeth gas flow was set to 20, and an auxiliary gas flow of 5 was used. Samples were analyzed in a randomized fashion bracketed by a blank and QC sample for background correction and normalization of the data, respectively. QC samples were additionally measured in confirmation and discovery mode to obtain further MS/MS spectra for identification. Obtained data sets have been processed by compound discoverer (CD) 3.1.0.035 (Thermo Fisher Scientific). Detailed description of the CD nodes and parameters can be found in the supporting information (**Supporting Figure S1**) and in the protocols section of the project MTBLS1782 in MetaboLights^11^. Compounds were annotated through comparing the retention time against our internal mass list database which was generated with authentic standard solutions. A retention time window of 0.4 min and a mass accuracy of 5 ppm for precursor masses was set. In addition, MS2 spectra were compared against our internal MS/MS database taking 10 ppm for fragment ion masses into account as well. HILIC-MS and RP-MS raw data file can be found here: **www.ebi.ac.uk/metabolights/MTBLS1782**.

Skyline (version 20.1.0) was used to create MS1 area based quality control charts^29^. The details of the targeted workflow for the specific metabolite classes of lipids, coenzymes, and carnitines can be found in the **supporting information** (**material and methods section**, *Targeted metabolomics of interesting metabolite classes*).

## Results and Discussion

Over the recent past, yeast-based standards have seen a slow but increasing acceptance in the metabolomics community. Several laboratories resorted to *Pichia pastoris* ethanolic yeast extracts^30–39^ primarily for ^13^C internal standardization. Only a few studies were related to instrument performance tests or method development in accordance to the here proposed application^33,38–40^.

### Metabolites and lipids present in the yeast material

The yeast extract was produced from controlled *Pichia pastoris* (Guillierm.) Phaff 1956 (*Komagataella phaffii Kurtzman*)^27^ fermentation followed by quenching and boiling ethanol metabolite extraction. One aliquot of yeast ethanolic extract contains 15 mg of dried cell extract originating from 20 mg yeast cell dry weight of *approx*. 2 billion cells. The metabolite inventory provided in this work is primarily based on data collected in two different laboratories (Köllensperger lab, University Vienna; Metabolomics Core Facility, VBCF, Vienna) using Orbitrap based HRMS workflows. The library assessment comprised orthogonal column chemistries which is reversed phase-liquid chromatography (RP-LC) and hydrophilic interaction liquid chromatographic (HILIC) separation at different pH values using positive and negative electrospray ionization. Typically, these workflows enable the annotation of metabolites at concentrations ranging from sub-nM to µM. Compounds identification was based on accurate mass (+/− 5 ppm), matching retention times, and MS2 spectra as compared to authentic standards and internal databases built upon these standards. While the major body of the library was obtained upon analysis of an aqueous dilution by the streamlined combination of HILIC-HRMS (at pH=8-9) and RP-LC-HRMS (at pH=2) (**raw data MTBLS1782**), some compounds such as lipids, carnitines, and coenzymes required alternative sample reconstitution/preparation and dedicated chromatographic separation (**Supporting Information**, material and methods part, *Targeted metabolomics of interesting metabolite classes*). More specifically, for coenzyme and acyl-carnitines analysis, a dedicated RP-LC-HRMS method utilizing ammonium bicarbonate at neutral pH^41^ was implemented. Coenzyme A, acetyl coenzyme A, L-carnitine, O-acetyl-L-carnitine, palmitoyl-L-carnitine, and propionyl-L-carnitine could be annotated. For lipid analysis, the dry ethanolic extract was reconstituted in 50% ACN and measured using RP-LC-HRMS with a gradient involving IPA. As a result, 27 consistently (different batch, different instruments) identified (accurate mass, retention time, and MS/MS comparison to a multi-lipid standard mix) phospholipids from the classes PC, PE, PG and PS (LPC 16:0, LPC 18:0, LPC 18:1, PC 34:0, PC 34:1, PC 34:2, PC 34:3, PC 34:4, PC 36:2, PC 36:3, PC 36:4, PC 36:5, PC 36:6, PE 34:1, PE 34:2, PE 34:3, PE 36:2, PE 36:3, PE 36:4, PE 36:5, PG 34:0, PG 36:0, PS 34:1, PS 34:2, PS 34:3, PS 36:2, PS 36:3) were recovered.

Literature search confirmed our findings and additionally expanded the reported compounds^30– 39^ in our library which can be found in the **Supplementary Table S1** (Excel sheet FinalMetaboliteList). As yeast is an eukaryotic organism, key regulatory elements are highly conserved between yeast and humans^19^. The curated list includes 206 metabolites and lipids (expected, detected, quantified in HMDB) present in the ethanolic yeast extract using orthogonal RP-LC-HRMS and HILIC-HRMS workflows of which 181 metabolites were reproducibly identified (at least in two different fermentation batches, different instruments, two participating laboratories or others^30–39^). The remaining 25 metabolites were reported elsewhere^30–33,36,37^(**Supplementary Table S1**, column, FinalMetaboliteList, “Additional literature hits”). **Figure 1** shows that the 206 metabolites represent a huge range of different metabolites from the super classes of (1) organic acids and derivatives making up 27% and the highest number of IDs in the presented metabolite followed by (2) nulceosides, nucleotides, and analogues, (3) lipids and lipid-like molecules, (4) organic oxygen compounds, (5) organoheterocyclic compounds, (6) organic nitrogen compounds and (7) benzoids. The classification in **Figure 1** is based on the classyFirer system^42^ categorizing in kingdoms, super classes, classes, sub classes, and molecular frameworks.

**Figure 1:**
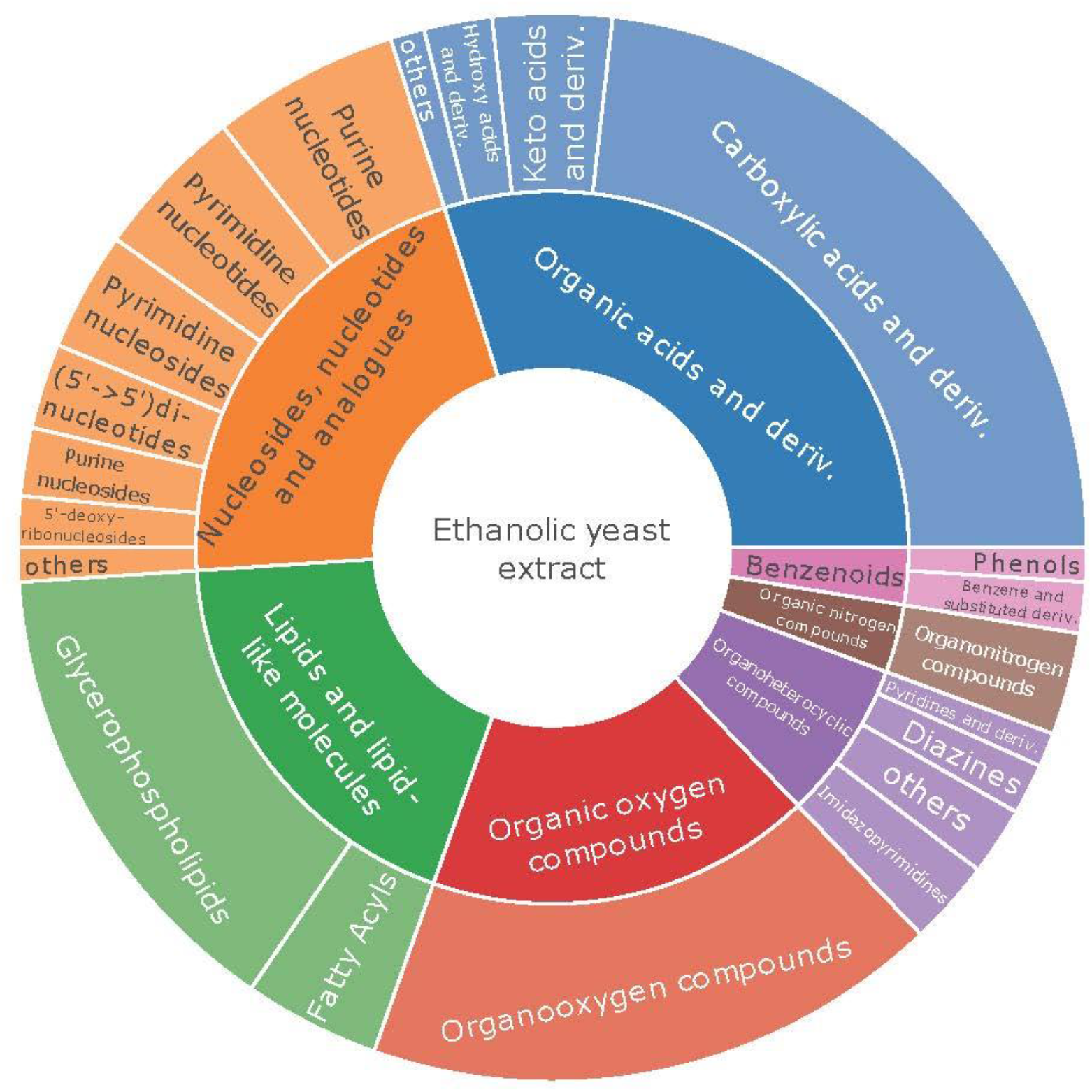
Annotated metabolite classes in ethanolic yeast extract using the classyFirer^42^ annotation system.

The human metabolome database (HMDB)^43,44^ was interrogated regarding overlaps with the yeast metabolome data base which is based on the Baker’s/Brewer’s yeast *Saccharomyces cerevisiae*^45^ crosschecking our library for plausibility. Most metabolites were present in the YMDB^45^ (170 from 206) and all metabolites were present in the HDMB^44^ (**Supplementary Table S1**). The higher match of our internal *Pichia pastoris* yeast extract database with the HMDB compared to YMDB can be explained by a more frequent comprehensive database curation of the HMDB (e.g: additional “status” ontology in HMDB including expected metabolites) and by the fact that the YMDB is based on the different yeast strain *S. cerevisiae*. The high coverage of the HDMB and the YMDB as well as the presence of several human key pathways proofs the potential of the proposed benchmark material. The related pathways playing a critical role in health and diseases are provided in the **Supplementary Table S1** (FinalMetaboliteList, column “pathway_names”). 100 metabolites were reported to be involved in cancer such as glucose (D-glucose, 6-phosphogluconic acid, gluconic acid, N-acetyl-D-glucosamine) and amino acids (L-arginine, L-asparagine, L-aspartic acid, L-cysteine, L-cystine, L-glutamic acid, L-glutamine, L-histidine, L-isoleucine, L-lactic acid, L-leucine, L-lysine, L-malic acid, L-methionine, L-phenylalanine, L-proline, L-serine, L-threonine, L-tryptophan, L-tyrosine, L-valine) which are uptake related compounds. These are known to be hallmarks of cancer metabolism^46^. Another example are obesity related pathways with a range of involved lipids (LysoPC(18:0), LysoPC(18:1). PC(16:0_18:1), PC(16:0_18:2), PC(16:0_18:3), PC(16:0_20:4), PC(18:0_18:3), PC(18:2_18:3), PC(18:3_16:1), PC(18:3_18:3)) and lipid precursors (L-carnitine, L-acetylcarnitine, L-palmitoylcarnitine, propionyl-L-carnitine), known to trigger obesity formation upon general lipid accumulation^47^.

Compared to a recent study on large scale metabolomics using three different human plasma reference materials similar classes and metabolite numbers (ca. 200) are covered^12^. However, such plasma reference materials are expensive, have to be pooled from several hundred individuals, and characterized prior to their use as quality control. Hence, they cannot be produced in the exact same way as it is possible using a stable fermentor and the same yeast strain. The general availability of the yeast material enables us to further expand the library on a regular basis by identifying metabolites with new commercial standards. It must be emphasized that the given *Pichia pastoris* inventory is restricted to the implemented biomass preparation resulting from fermentation conditions, the number of cells submitted to extraction, and the ethanolic extraction procedure itself. The panel can be significantly changed upon using different extractions. The use of methanol/chloroform (Folch) extraction^48^ for example expanded the lipid panel with regard to non-polar lipids such as triglycerides, diglycerides, ceramides or ergosterol^21,49,50^.

Overall, *Pichia pastoris* yeast extract is an excellent source to perform quality controls in non-targeted metabolomics experiments as reproducible and cheap production in high amounts is possible yielding an interesting panel of essential eukaryotic metabolites and lipids.

### Yeast quality control for instrument performance

Benchmarking can be performed at different levels, either for checking instrumental/method performance in general (e.g. upon introduction of new MS instrumentation) or for daily quality control. In principle, the complete library is available for performance tests provided that the yeast derived standard is stable under defined storage conditions. In order to proof the suitability of the proposed benchmark, NTA by RP-LC/HILIC-HRMS was performed in ethanolic extracts from different fermentation batches (3 fermentations performed in 2017, 2018, 2019, respectively), which had been stored at −80 °C. The streamlined workflow (HILIC at pH 8, RP-LC at pH 2) obtained from one laboratory (Metabolomics Core Facility, VBCF Vienna) within the same sequence revealed a list of 151 compounds (see **Supplementary Table S1**, Batch comparison). After filtering for stable compounds (response ratios between 0.5 and 5 was set as criterion), 126 (pHILIC-MS: 76 metabolites, RP-MS: 50 metabolites) were obtained leading to 80 unique (46 metabolites were identified in both HILIC and RP) compounds (**Supplementary Table S1**, BC filtered ratios 0.5 to 5). This restricted list of annotated compounds was present in all three fermentation batches and the pooled control. The variation of each metabolite within one batch group was also excellent and is reflected by the group coefficient variant (CV) which was on average 11%. A low concentration of a metabolite might be responsible for elevated group CVs and compounds with elevated group CVs through inappropriate integration of the software are marked with an asterisk.

Another way of visualizing sample comparability are box-and-whiskers plots showing data distribution **(Supplement Figure S2. A** and **S2. B)** automatically generated by CD. Each box corresponds to annotated metabolite areas on a logarithmic scale of each sample measured with RP and HILIC, respectively. The median value of the dataset is represented through a line in the rectangle. Outlier data points with either very high or very low metabolite area lie above or below the respective whiskers. The relatively broad concentration range of the panel of observed metabolites in yeast is responsible for the high variation. Again, good repeatability through all yeast fermentation samples and quality controls is observed:).

Thus, the obtained biological repeatability of the fully controlled fermentations together with the storage stability of the dried extraction aliquots over three years, makes the yeast benchmark fit for purpose for a whole range of metabolomics measurement platforms. **Figure 2 A + B** show the application of the yeast benchmark as our in-house quality check routine of a lipidomics method. 26 lipids were analyzed by RP-LC-HRMS (LPC 16:0, LPC 18:0, PC 34:0, PC 34:1, PC 34:2, PC 34:3, PC 34:4, PC 36:2, PC 36:3, PC 36:4, PC 36:5, PC 36:6, PE 34:1, PE 34:2, PE 34:3, PE 36:2, PE 36:3, PE 36:4, PE 36:5, PG 34:0, PG 36:0, PS 34:1, PS 34:2, PS 34:3, PS 36:2, PS 36:3) on two instruments (Q Exactive™ HF Hybrid Quadrupole-Orbitraps, instrument HF 1 and HF 2) and are plotted with respect to peak area and retention time in order to control the instrumental performance prior measurement sequences. The measurement from January 27 2020 (200127_HF2) showed significant signal decrease requiring cleaning of the instrument (S-lens, shield cleaning). The signal improved as validated with the in-house yeast lipid quality control from February 2020 (**Figure 2.B**, 2002010_HF2_1 and 20200210_HF2_2). Not considering measurement outliers (i.e. 200127_HF2 replicate 1 and 2), the retention time repeatability over the course of 10 months for each instrument was excellent with most of the lipids (19 out of 26 lipids) having a variation < 5% (ranging from 0.4% for high abundant PE 34:1 with areas of 10^6^ to 10% for low abundant PI 34:3 with areas of 10^3^). **Figure 2.A** shows all lipids while **Supporting Figure S3** depicts the biggest lipid class of PC for further insight. The overall plotted peak areas cover 5 orders of magnitudes, resembling the typical situation in metabolomics type of measurements (**Figure 2.B**).

**Figure 2.A:**
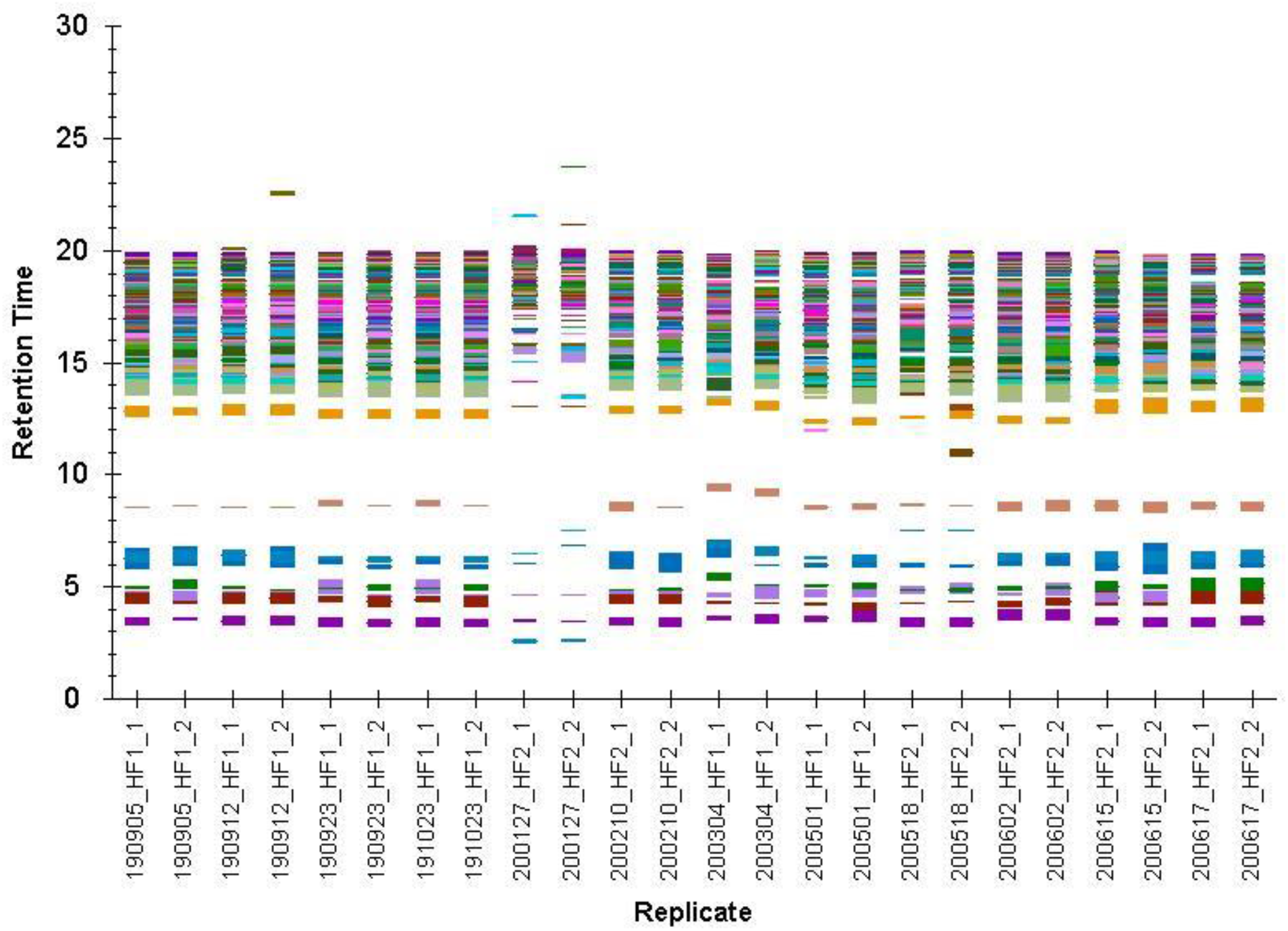
Yeast extract retention time stability of the RP-HRMS targeted lipid method on two different high-resolution mass spectrometers (Orbitrap HF 1, HF 2) over the time course of 10 months using Skyline for visualization^29^.

**Figure 2.B.:**
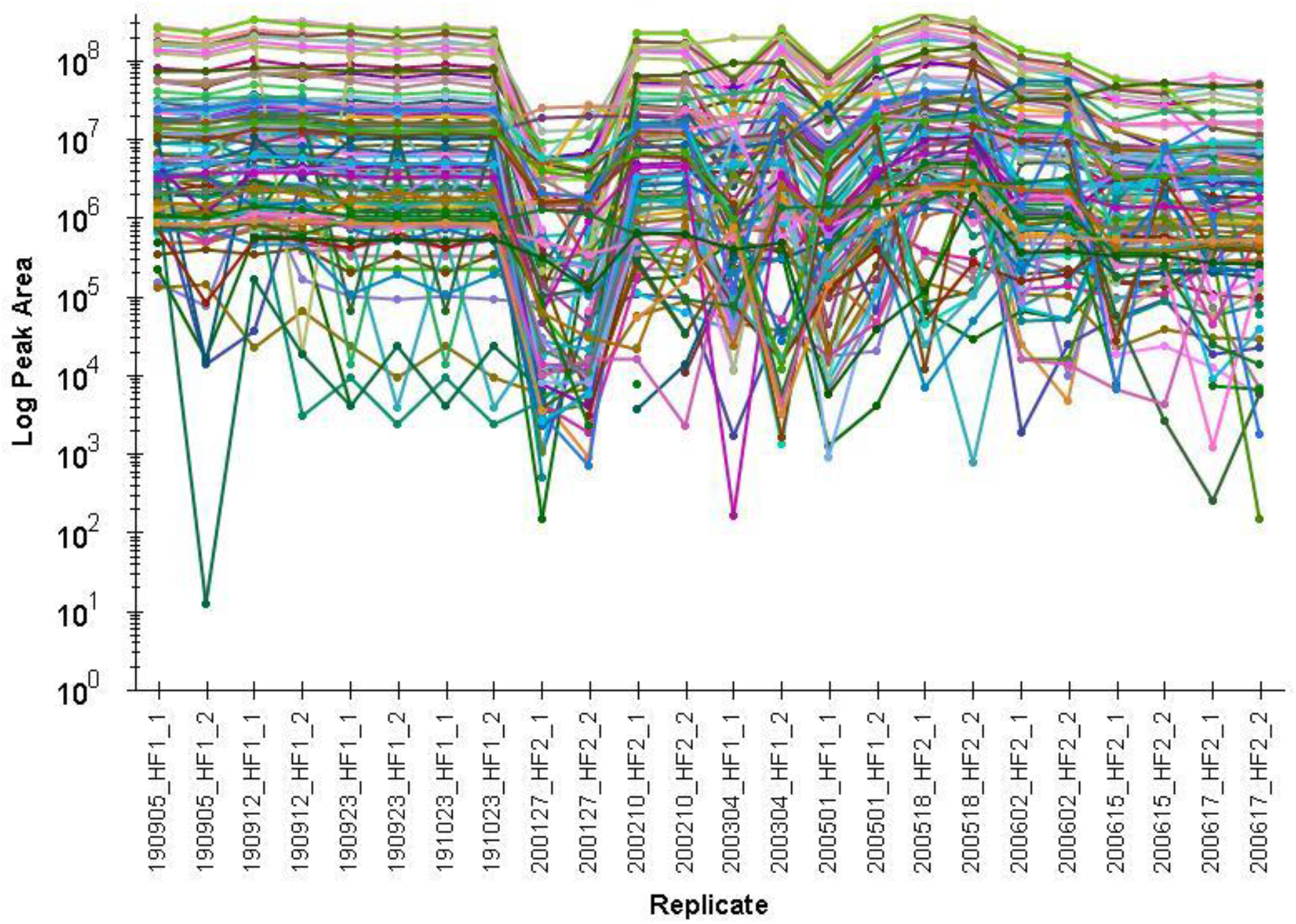
Yeast extract control of 26 lipids using RP-HRMS to monitor the performance of two Orbitrap mass analyzers (HF 1, HF 2) using Skyline for visualization^29^.

### Yeast quality controls facilitating method development

No single method or platform can cover the full metabolome due to the polarity range of metabolites (from highly water-soluble sugars to very lipophilic triacylglycerols)^51^. Benchmarking materials by providing a defined panel of metabolites in biological matrix can facilitate the development and proper validation of mass spectrometry-based assays. A metric is introduced allowing to evaluate the metabolome coverage with regard to other platforms/procedures. In our laboratory, we successfully applied endogenous and labeled yeast extracts for method validation of different LC-MS based workflows such as (1) merged metabolomics and lipidomics workflow based on the combination of HILIC and RP^52^, (2) anion exchange chromatography coupled to high-resolution MS^53^ (3) polar and non-polar lipid analysis by online HILIC and RP combination^54^ and (4) comparison of different LC-MS based workflows^40,55^. In this work, high metabolite coverage of ethanolic yeast extracts was achieved using orthogonal HILIC and RP-MS (**Supplementary Table S1**, Batch comparison). To extend the metabolome coverage of the benchmarking material, additional LC-MS methods were developed to target (1) acylcarnitines and coenzymes (lower abundance and problems with ionization efficiency using standard metabolomics workflows) and (2) lipids (different solubility). Using the targeted neutral RP method, coenzyme A, acetyl coenzyme A, L-carnitine, O-acetyl-L-carnitine, palmitoyl-L-carnitine, and propionyl-L-carnitine were identified (MS2 matching to commercial standards, m/z, RT; exemplary shown for O-acetyl-L-carnitine, **Supporting Figure S4**) and led to estimated concentrations (comparison to 5 µM standard mix) in the low to high nM range (**Supporting Figure S5**). The dedicated lipid analysis led to the identification of 26 additional phospholipids as described above to embrace the broad diversity of the metabolome (**Figure 2.A**).

These results show the high metabolome coverage of the yeast material which can be interrogated to develop dedicated LC-MS methods for specific metabolite classes. Therefore, yeast ethanolic extracts are an ideal test matrix to establish new metabolomics workflows.

## Conclusion

In the last years, yeast ethanolic extracts were extensively used in our laboratory produced by the described controlled *Pichia pastoris* fermentation. We found that the material provides a sufficiently large coverage of the eukaryotic metabolome. Our reproducible metabolite database derived from different fermentation batches which were measured in two different laboratories. Additional metabolites were incorporated into the library that were identified in the literature and measured by groups working with this yeast material. The provided list of 206 metabolites covers the classes of 1) organic acids and derivatives (2) nulceosides, nucleotides, and analogues, (3) lipids and lipid-like molecules, (4) organic oxygen compounds, (5) organoheterocyclic compounds, (6) organic nitrogen compounds and (7) benzoids and can be further extended using commercially available standards and different extraction strategies. All yeast metabolites were also reported in the human metabolome database with most of the metabolites being stable for several years in comparable concentrations.

Commercially available materials such as human plasma reference materials (e.g. SRM 1950 or CHEAR) are expensive. As they have to be pooled from several hundred individuals and characterized prior to their use as quality control, reference materials from human origin can never be reproduced in the exact same way. The yeast ethanolic extract is of eukaryotic origin, easily accessible, and can be produced under controlled fermentation condition. Hence, the yeast material is an ideal starting point for broader community-wide used NTA quality controls, similar to HeLa cells in proteomics. Expansion of the database on a regular basis in our laboratory as well as extensive use by other metabolomics laboratories will lead to a deep characterization of these extracts and a very detailed list of the contained metabolites. Moreover, batch normalization is feasible with ethanolic yeast extracts. Hence, we dare to propose the described yeast material as an ideal open-source database to benchmark the coverage of non-targeted metabolomics workflows.

## Acknowledgement

This work was supported by the University of Vienna, the Faculty of Chemistry, the Vienna Metabolomics Center (VIME; http://metabolomics.univie.ac.at/), and the research platform Chemistry Meets Microbiology of the University of Vienna. The authors thank Martin Schaier for graphical abstract preparation and all members of the Environmental Analysis group (University of Vienna) for continuous support. The service untargeted metabolomics was performed at the VBCF Metabolomics unit (www.vbcf.ac.at) and funded by the City of Vienna through the Vienna Business Agency.

## Supporting information

**Supplementary Table 1:** (1) list of metabolites and lipids identified in ethanol extract, (2) batch comparison (2017/2018/2019), (3) group area batch comparison ratios filtered

### Extended methods section and additional information on materials and methods

**S1**. Compound Discoverer 3.1.0.305 workflow

**S2**. Box plots of annotated metabolite areas of each sample on a logarithmic scale measured with RP-HRMS and HILIC-HRMS and derived from compound discoverer

**S3**. Retention time of the yeast control stability in the exemplary class of PCs

**S4**. MS2 of O-Acetyl-L-carnitine in 5 µM standard and ethanolic yeast extracts

**S5**. Estimated carnitine and coenzyme concentrations in ethanolic yeast extract

## Author Contributions

^**#**^ The manuscript was written through contributions of all authors. All authors have given approval to the final version.

## Notes

The authors declare the following competing financial interest: G.H. is working for the University Vienna spin-off of Isotopic Solutions producing endogenous and labeled *Pichia pastoris* yeast extract.

## For TOC only

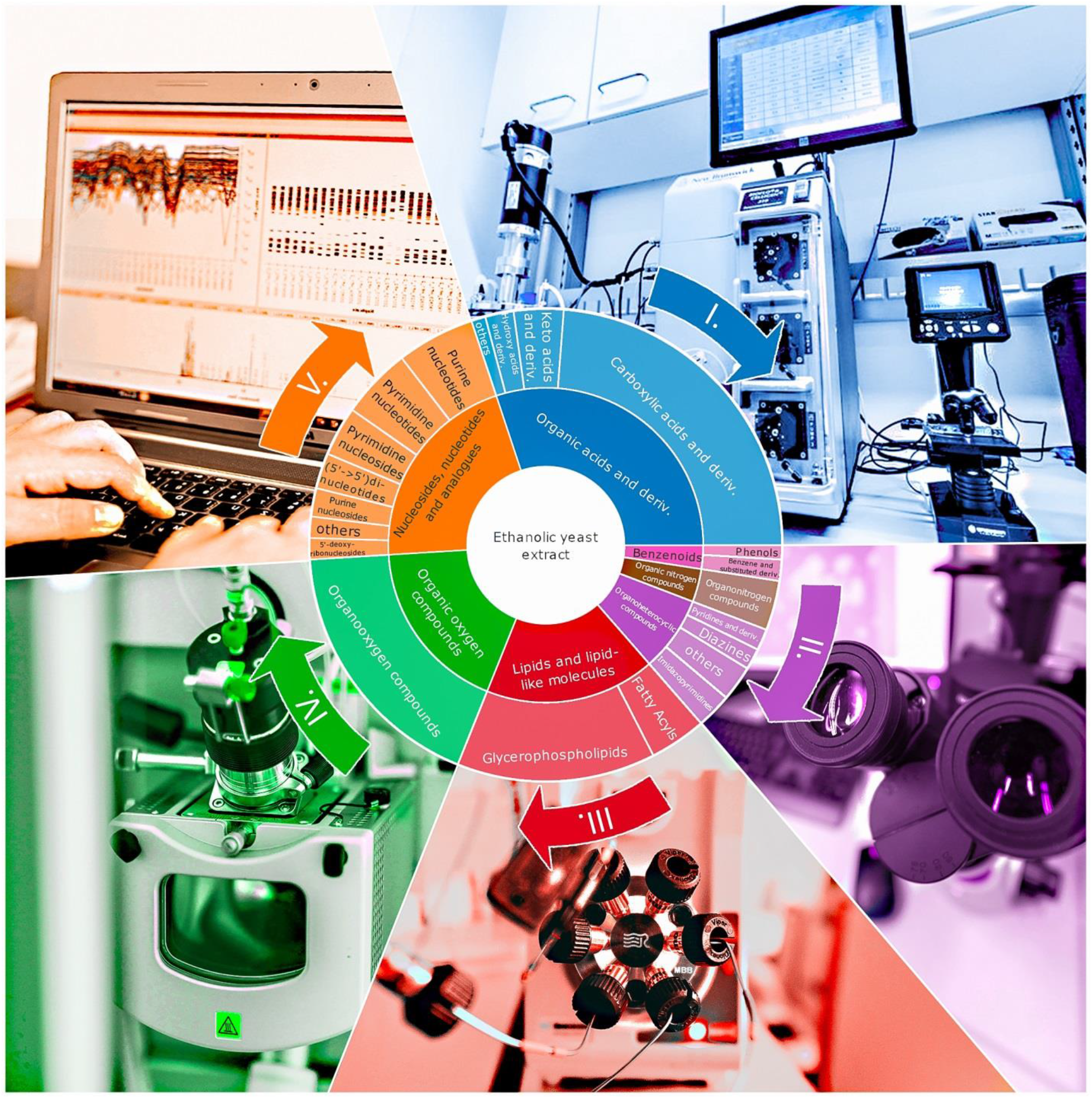

## References

(1) Baker, M. Nat. Methods 2011, 8 (2), 117–121.

(2) Schymanski, E. L.; Singer, H. P.; Slobodnik, J.; Ipolyi, I. M.; Oswald, P.; Krauss, M.; Schulze, T.; Haglund, P.; Letzel, T.; Grosse, S.; Thomaidis, N. S.; Bletsou, A.; Zwiener, C.; Ibáñez, M.; Portolés, T.; De Boer, R.; Reid, M. J.; Onghena, M.; Kunkel, U.; Schulz, W.; Guillon, A.; Noyon, N.; Leroy, G.; Bados, P.; Bogialli, S.; Stipaničev, D.; Rostkowski, P.; Hollender, J. Anal. Bioanal. Chem. 2015, 407 (21), 6237–6255.

(3) Pinu, F. R. Food Res. Int. 2015, 72, 80–81.

(4) Naz, S.; Vallejo, M.; García, A.; Barbas, C. J. Chromatogr. A 2014, 1353, 99–105.

(5) Dudzik, D.; Barbas-Bernardos, C.; García, A.; Barbas, C. J. Pharm. Biomed. Anal. 2018, 147, 149–173.

(6) González-Riano, C.; Dudzik, D.; Garcia, A.; Gil-de-la-Fuente, A.; Gradillas, A.; Godzien, J.; López-Gonzálvez, Á., Rey-Stolle, F.; Rojo, D.; Ruperez, F. J.; Saiz, J.; Barbas, C. Anal. Chem. 2020, 92 (1), 2047–217X-2–13.

(7) Metabolomics Society: Standardization in Metabolomics Experiments http://metabolomicssociety.org/resources/metabolomics-standards (accessed Jul 14, 2020).

(8) MSI Board Members Sansone, S.-A., Fan, T.; Goodacre, R.; Griffin, J. L.; Hardy, N. W.; Kaddurah-Daouk, R.; Kristal, B. S.; Lindon, J.; Mendes, P.; Morrison, N.; Nikolau, B.; Robertson, D.; Sumner, L. W.; Taylor, C.; van der Werf, M.; van Ommen, B.; Fiehn, O. Nat. Biotechnol. 2007, 25 (8), 846–848.

(9) Sumner, L. W.; Samuel, T.; Noble, R.; Gmbh, S. D.; Barrett, D.; Beale, M. H.; Hardy, N. Metabolomics 2007, 3 (3), 211–221.

(10) Salek, R. M.; Steinbeck, C.; Viant, M. R.; Goodacre, R.; Dunn, W. B. Gigascience 2013, 2 (1), 2047-217X-2–13.

(11) Haug, K.; Cochrane, K.; Nainala, V. C.; Williams, M.; Chang, J.; Jayaseelan, K. V.; O’Donovan, C. Nucleic Acids Res. 2020, 48 (D1), D440–D444.

(12) Liu, K. H.; Nellis, M.; Uppal, K.; Ma, C.; Tran, V.; Liang, Y.; Walker, D. I.; Jones, D. P. Anal. Chem. 2020, 92 (13), 8836–8844.

(13) Altelaar, A. F. M.; Frese, C. K.; Preisinger, C.; Hennrich, M. L.; Schram, A. W.; Timmers, H. T. M.; Heck, A. J. R.; Mohammed, S. J. Proteomics 2013, 88, 14–26.

(14) Ahn, N. G.; Shabb, J. B.; Old, W. M.; Resing, K. A. ACS Chem. Biol. 2007, 2 (1), 39–52.

(15) Navarro, P.; Kuharev, J.; Gillet, L. C.; Bernhardt, O. M.; MacLean, B.; Röst, H. L.; Tate, S. A.; Tsou, C. C.; Reiter, L.; Distler, U.; Rosenberger, G.; Perez-Riverol, Y.; Nesvizhskii, A. I.; Aebersold, R.; Tenzer, S. Nat. Biotechnol. 2016, 34 (11), 1130–1136.

(16) Kelstrup, C. D.; Bekker-Jensen, D. B.; Arrey, T. N.; Hogrebe, A.; Harder, A.; Olsen, J. V. J. Proteome Res. 2018, 17 (1), 727–738.

(17) Köcher, T.; Pichler, P.; Swart, R.; Mechtler, K. Nat. Protoc. 2012, 7 (5), 882–890.

(18) Ridgeway, M. E.; Bleiholder, C.; Mann, M.; Park, M. A. TrAC – Trends Anal. Chem. 2019, 116, 324–331.

(19) Nielsen, J. FEBS Lett. 2009, 583 (24), 3905–3913.

(20) Schwaiger, M.; Rampler, E.; Hermann, G.; Miklos, W.; Berger, W.; Koellensperger, G.; Anal. Chem. 2017, 89 (14), 7667–7674.

(21) Rampler, E.; Coman, C.; Hermann, G.; Sickmann, A.; Ahrends, R.; Koellensperger, G. Analyst 2017, 142 (11), 1891–1899.

(22) Neubauer, S.; Haberhauer-Troyer, C.; Klavins, K.; Russmayer, H.; Steiger, M. G.; Gasser, B.; Sauer, M.; Mattanovich, D.; Hann, S.; Koellensperger, G. J. Sep. Sci. 2012, 35 (22), 3091–3105.

(23) Hermann, G.; Schwaiger, M.; Volejnik, P.; Koellensperger, G. J. Pharm. Biomed. Anal. 2018, 155, 329–334.

(24) Schwaiger, M.; Schoeny, H.; El Abiead, Y.; Hermann, G.; Rampler, E.; Koellensperger, G. Analyst 2019, 144 (1), 220–229.

(25) Rampler, E.; Criscuolo, A.; Zeller, M.; El Abiead, Y.; Schoeny, H.; Hermann, G.; Sokol, E.; Cook, K.; Peake, D. A.; Delanghe, B.; Koellensperger, G. Anal. Chem. 2018, 90 (11), 6494–6501.

(26) Guijas, C.; Montenegro-Burke, J. R.; Domingo-Almenara, X.; Palermo, A.; Warth, B.; Hermann, G.; Koellensperger, G.; Huan, T.; Uritboonthai, W.; Aisporna, A. E.; Wolan, D. W.; Spilker, M. E.; Benton, H. P.; Siuzdak, G. Anal. Chem. 2018, 90 (5), 3156–3164.

(27) Kurtzman, C. P. J. Ind. Microbiol. Biotechnol. 2009, 36 (11), 1435–1438.

(28) Wernisch, S.; Pennathur, S. Anal. Bioanal. Chem. 2016, 408 (22), 6079–6091.

(29) Adams, K. J.; Pratt, B.; Bose, N.; Dubois, L. G.; St. John-Williams, L., Perrott, K. M.; Ky, K.; Kapahi, P.; Sharma, V.; Maccoss, M. J.; Moseley, M. A.; Colton, C. A.; Maclean, B. X.; Schilling, B.; Thompson, J. W. J. Proteome Res. 2020, 19 (4), 1447–1458.

(30) Si-Hung, L.; Troyer, C.; Causon, T.; Hann, S. Talanta 2019, 205, 120147.

(31) Mairinger, T.; Sanderson, J.; Hann, S. Anal. Bioanal. Chem. 2019, 411 (8), 1495–1502.

(32) Demarest, T. G.; Truong, G. T. D.; Lovett, J.; Mohanty, J. G.; Mattison, J. A.; Mattson, M. P.; Ferrucci, L.; Bohr, V. A.; Moaddel, R. Anal. Biochem. 2019, 572, 1–8.

(33) Si-Hung, L.; Causon, T. J.; Hann, S. Electrophoresis 2017, 38 (18), 2287–2295.

(34) Galvez, L.; Rusz, M.; Schwaiger-Haber, M.; El Abiead, Y.; Hermann, G.; Jungwirth, U.; Berger, W.; Keppler, B. K.; Jakupec, M. A.; Koellensperger, G. Metallomics 2019, 11 (10), 1716–1728.

(35) Mairinger, T.; Weiner, M.; Hann, S.; Troyer, C. Anal. Chem. 2020, 92 (7), 4875–4883.

(36) Puleston, D. J.; Buck, M. D.; Klein Geltink, R. I., Kyle, R. L.; Caputa, G.; O’Sullivan, D.; Cameron, A. M.; Castoldi, A.; Musa, Y.; Kabat, A. M.; Zhang, Y.; Flachsmann, L. J.; Field, C. S.; Patterson, A. E.; Scherer, S.; Alfei, F.; Baixauli, F.; Austin, S. K.; Kelly, B.; Matsushita, M.; Curtis, J. D.; Grzes, K. M.; Villa, M.; Corrado, M.; Sanin, D. E.; Qiu, J.; Pällman, N.; Paz, K.; Maccari, M. E.; Blazar, B. R.; Mittler, G.; Buescher, J. M.; Zehn, D.; Rospert, S.; Pearce, E. J.; Balabanov, S.; Pearce, E. L. Cell Metab. 2019, 30 (2), 352–363.

(37) Swain, A.; Bambouskova, M.; Kim, H.; Andhey, P. S.; Duncan, D.; Auclair, K.; Chubukov, V.; Simons, D. M.; Roddy, T. P.; Stewart, K. M.; Artyomov, M. N. Nat. Metab. 2020.

(38) Chu, D. B.; Troyer, C.; Mairinger, T.; Ortmayr, K.; Neubauer, S.; Koellensperger, G.; Hann, S. Anal. Bioanal. Chem. 2015, 407 (10), 2865–2875.

(39) Mairinger, T.; Hann, S. Anal. Bioanal. Chem. 2017, 409 (15), 3713–3718.

(40) Schwaiger-Haber, M.; Hermann, G.; El Abiead, Y.; Rampler, E.; Wernisch, S.; Sas, K.; Pennathur, S.; Koellensperger, G. Anal. Bioanal. Chem. 2019, 411 (14), 3103–3113.

(41) Neubauer, S.; Chu, D. B.; Marx, H.; Sauer, M.; Hann, S.; Koellensperger, G. Anal. Bioanal. Chem. 2015, 407 (22), 6681–6688.

(42) Djoumbou Feunang, Y., Eisner, R.; Knox, C.; Chepelev, L.; Hastings, J.; Owen, G.; Fahy, E.; Steinbeck, C.; Subramanian, S.; Bolton, E.; Greiner, R.; Wishart, D. S. J. Cheminform. 2016, 8 (1), 1–20.

(43) Wishart, D. S.; Knox, C.; Guo, A. C.; Eisner, R.; Young, N.; Gautam, B.; Hau, D. D.; Psychogios, N.; Dong, E.; Bouatra, S.; Mandal, R.; Sinelnikov, I.; Xia, J.; Jia, L.; Cruz, J. A.; Lim, E.; Sobsey, C. A.; Shrivastava, S.; Huang, P.; Liu, P.; Fang, L.; Peng, J.; Fradette, R.; Cheng, D.; Tzur, D.; Clements, M.; Lewis, A.; de souza, A.; Zuniga, A.; Dawe, M.; Xiong, Y.; Clive, D.; Greiner, R.; Nazyrova, A.; Shaykhutdinov, R.; Li, L.; Vogel, H. J.; Forsythei, I. Nucleic Acids Res. 2009, 37, 603–610.

(44) Wishart, D. S.; Jewison, T.; Guo, A. C.; Wilson, M.; Knox, C.; Liu, Y.; Djoumbou, Y.; Mandal, R.; Aziat, F.; Dong, E.; Bouatra, S.; Sinelnikov, I.; Arndt, D.; Xia, J.; Liu, P.; Yallou, F.; Bjorndahl, T.; Perez-Pineiro, R.; Eisner, R.; Allen, F.; Neveu, V.; Greiner, R.; Scalbert, A. Nucleic Acids Res. 2013, 41 (D1), 801–807.

(45) Ramirez-Gaona, M.; Marcu, A.; Pon, A.; Guo, A. C.; Sajed, T.; Wishart, N. A.; Karu, N.; Feunang, Y. D.; Arndt, D.; Wishart, D. S. Nucleic Acids Res. 2017, 45 (D1), D440–D445.

(46) Pavlova, N. N.; Thompson, C. B. Cell Metab. 2016, 23 (1), 27–47.

(47) Meikle, P. J.; Summers, S. A. Nat. Rev. Endocrinol. 2017, 13 (2), 79–91.

(48) Folch, J.; Lees, M.; Sloane Stanley, G. H. H. J. Biol. Chem 1952, 226, 497–509.

(49) Grillitsch, K.; Tarazona, P.; Klug, L.; Wriessnegger, T.; Zellnig, G.; Leitner, E.; Feussner, I.; Daum, G. Biochim. Biophys. Acta 2014, 1838 (7), 1889–1897.

(50) Klug, L.; Tarazona, P.; Gruber, C.; Grillitsch, K.; Gasser, B.; Trötzmüller, M.; Köfeler, H.; Leitner, E.; Feussner, I.; Mattanovich, D.; Altmann, F.; Daum, G. Biochim. Biophys. Acta – Mol. Cell Biol. Lipids 2014, 1841 (2), 215–226.

(51) Cajka, T.; Fiehn, O. Anal Chem 2016, 88, 524–545.

(52) Schwaiger, M.; Schoeny, H.; El Abiead, Y.; Hermann, G.; Rampler, E.; Koellensperger, G. Analyst 2019, 144 (1), 220–229.

(53) Schwaiger, M.; Rampler, E.; Hermann, G.; Miklos, W.; Berger, W.; Koellensperger, G. Anal. Chem. 2017, 89 (14), 7667–7674.

(54) Rampler, E.; Schoeny, H.; Mitic, B. M.; El Abiead, Y.; Schwaiger, M.; Koellensperger, G. Analyst 2018, 143 (5), 1250–1258.

(55) Rampler, E.; Schoeny, H.; Schwaiger-Haber, M.; Koellensperger, G. In Reference Module in Food Science; Elsevier, 2020.

