## Supplementary Information for "Benchmarking non-targeted metabolomics using yeast derived libraries"

**Supporting information**

This section contains the extended methods section and additional information on materials and methods, (S1) **compound Discoverer 3.1.0.305 workflow, (S2) RP-HRMS and HILIC-HRMS batch comparison of the metabolites areas derived from compound discoverer,** (S3) retention time stability of the yeast control in the exemplary class of PCs, **(S4)** MS2 of O-Acetyl-L-carnitine in 5 µM standard and ethanolic yeast extracts**, and (S5)** estimated carnitine and coenzyme concentrations in ethanolic yeast extract. Additionally, a (1) *list of metabolites and lipids identified in ethanol extract*, (2) *batch comparison (2017/2018/2019), and (3) group area batch comparison ratios filtered (0.5 to 5)* can be found in **Supplementary** **Table S1.**

**Material and Methods**

*Targeted metabolomics of interesting metabolite classes*

*Lipids*

The dried yeast extract was dissolved in 2 mL 50% acetontrile for lipid analysis using reversed-phase separation with an isopropanol gradient. The chromatographic separation was performed using an Acquity HSS T3 (2.1 mm × 150 mm, 1.8 μm, Waters) with a VanGuard pre-column. Solvent A was ACN/H_2_O (3:2, v/v), and solvent B was IPA/ACN (9:1, v/v), both solvents contained 0.1% formic acid, and 10 mM ammonium formate. The following gradient was applied: 0-2 min: 30% B, 2-15 min: ramp to 75% B, 15-17 min: to 100 % B, 17-22 min: holding 100% B, 22-27 min: 30% B. The injector needle was washed with 75% IPA, 24.9% H_2_O, and 0.1% formic acid prior to each injection. High-resolution MS with a Q Exactive HF (Thermo Fisher Scientific) was performed using ddMS2 (Top10) for lipid detection and the following HESI source parameters: capillary temperature of 270 °C, sheath gas flow rate of 50, auxiliary flow rate of 14, sweep gas of 3, S-lens RF level of 45 and auxiliary gas heater temperature of 380 °C applying a spray voltage of 3.5 kV in positive mode and 2.8 kV in negative mode.

*Carnitines and Coenzymes*

The dried yeast extract was dissolved in 2 mL 100% water for carnitines and coenzymes analysis by reversed-phase separation adapted from Neubauer et al^8^. The chromatographic separation was performed using an Acquity HSS T3 with pre-column (as stated for the lipids). Solvent A was 50 mM NH_4_HCO_3_, pH 6.94 and solvent B was 100% ACN and the following gradient was applied: 0-7 min: ramp from 100% A to 20% B, 7-8.1 min: ramp to 40% B, 8.1-10 min: 40% B, 10.1-13 min: 100% A. MS detection was performed by Q Exactive HF (Thermo Fisher Scientific) using positive ionization mode (+3.5 kV), S-lens RF level of 30 and ddMS2 (Top10) for carnitine and coenzyme analysis.

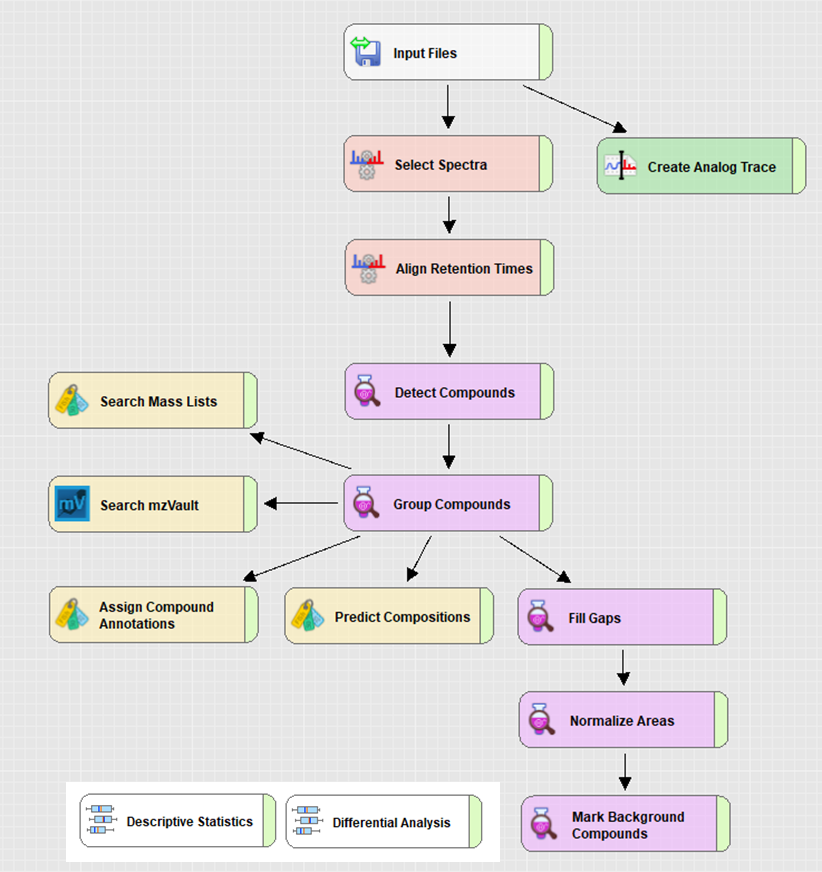

**Supporting Figure S1: Compound Discoverer 3.1.0.305 workflow for the untargeted RP and HILIC-MS analysis. Detailed parameters can be found here:** [www.ebi.ac.uk/metabolights/MTBLS1782](https://www.ebi.ac.uk/metabolights/editor/www.ebi.ac.uk/metabolights/MTBLS1782)

**Supporting Figure 2. A, Figure 2. B**

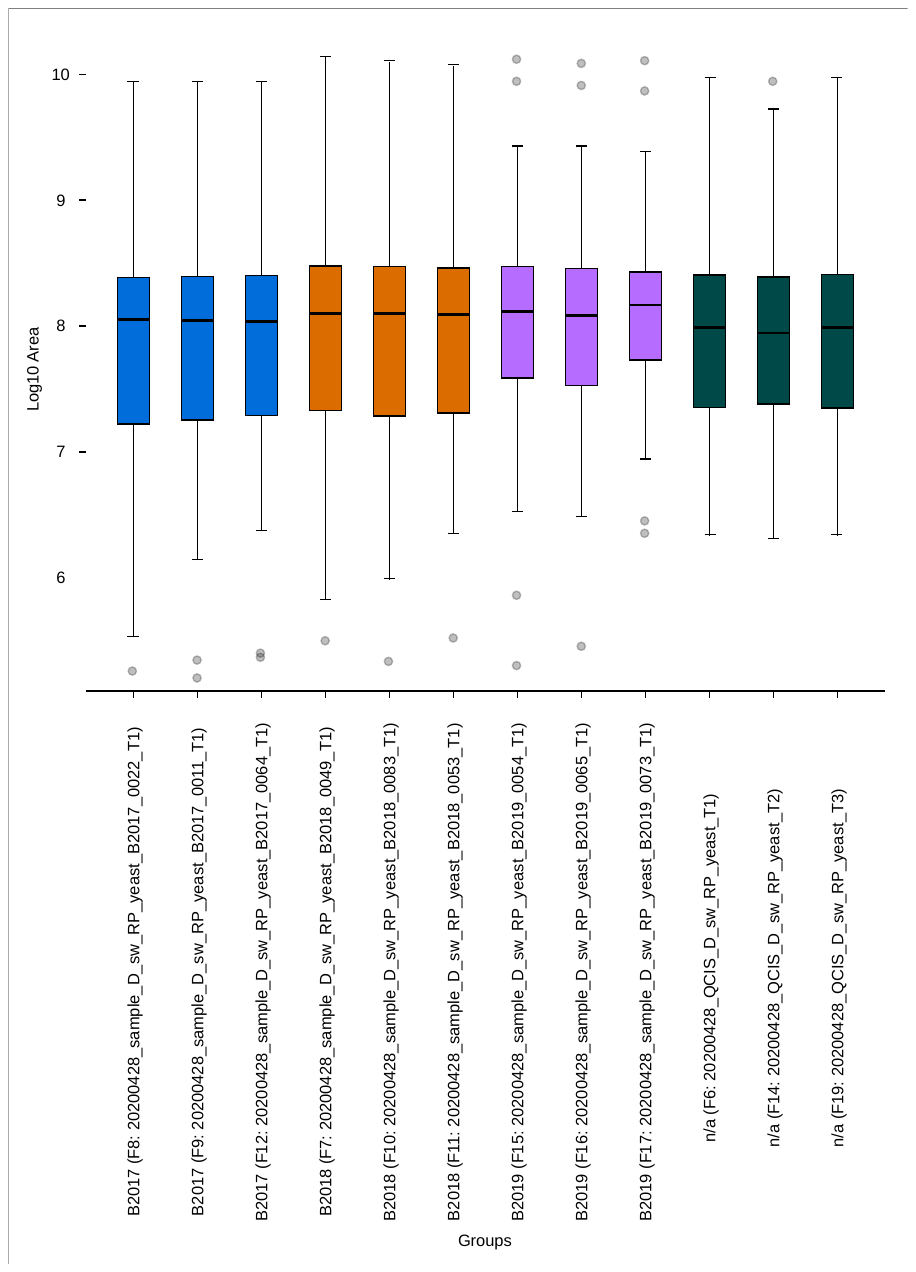

**Supporting Figure S2.A.** Box-and-whisker plot of the data distribution. Each box corresponds to the annotated metabolite areas of each sample on a logarithmic scale (log_10_) measured with RP-HRMS. Ethanolic yeast extracts (n=3) from three different years (2017, 2018, 2019) and the mixed quality control (n=3) are compared. Variance depicts the broad concentration range of the metabolome.

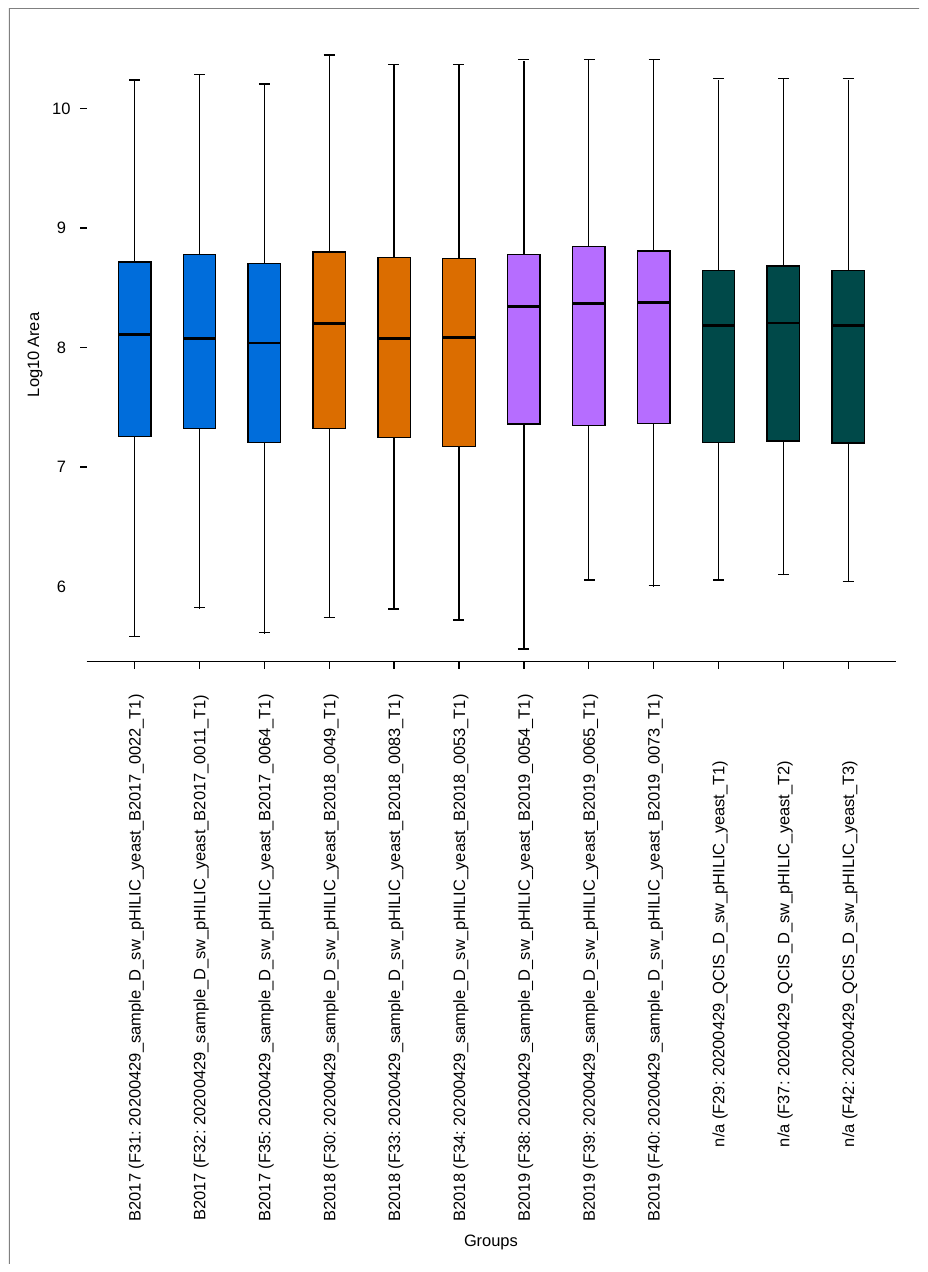

**Supporting Figure 2.B**. Box-and-whisker plot of the data distribution. Each box corresponds to the annotated metabolite areas of each sample on a logarithmic scale (log_10_) measured with HILIC-HRMS. Ethanolic yeast extracts (n=3) from three different years (2017, 2018, 2019) and the mixed quality control (n=3) are compared. Variance depicts the broad concentration range of the metabolome.

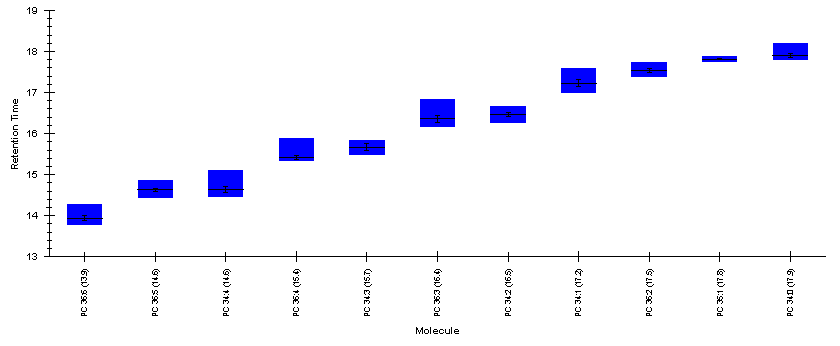

**Supporting Figure S3:** Retention time stability of the exemplary class of PCs in the yeast control followed over the time course of 10 months.

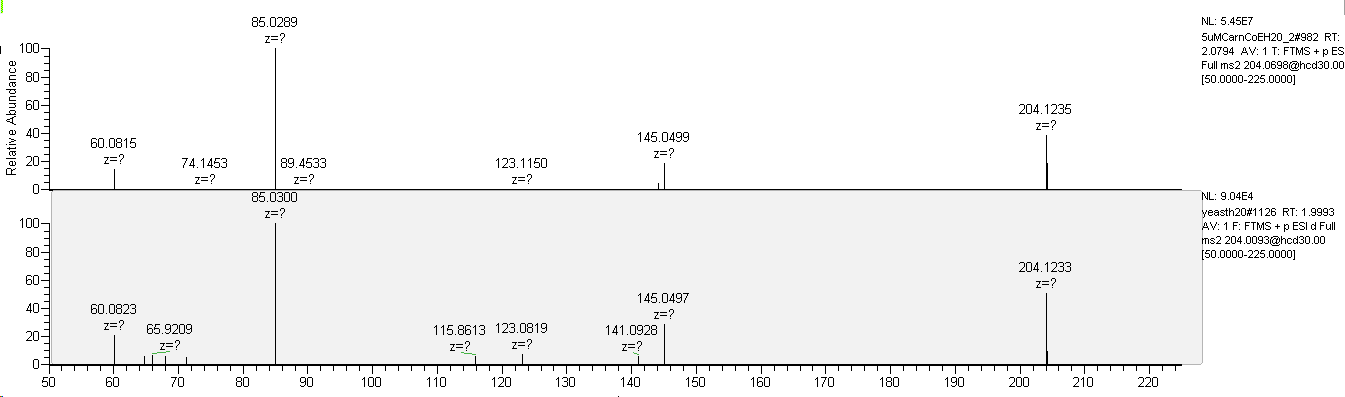

**Supporting Figure S4:** MS^2^ of O-Acetyl-L-carnitine in 5 µM standard and ethanolic yeast extracts measured by neutral RP-HRMS in positive mode.

**Supporting Table S5:** Estimated carnitine and coenzyme concentrations in ethanolic yeast extract determined using neutral RP-HRMS and 5 µM commercially available standards

|  |  | | **M+H** | | **RT** | | **Est. Concentration (nM)** |
| --- | --- | --- | --- | --- | --- | --- | --- |
| **Coenzyme A** | C_21_H_36_N_7_O_16_P_3_S | 768.1225 | | 6.29 | | 2792 | |
| **Acetyl coenzyme A** | C_23_H_38_N_7_O_17_P_3_S | 810.1330 | | 6.6 | | 316 | |
| **Palmitoyl coenzyme A** | C_37_H_66_N_7_O_17_P^3^S | 1006.3522 | | 7.1 | | < LOQ | |
| **Carnitine** | C_7_H_15_NO_3_ | 162.1125 | | 1.48 | | 139 | |
| **O-Acetyl-L-carnitine** | C_9_H_17_NO_4_ | 204.1230 | | 2.16 | | 16 | |
| **Propionyl-L-Carnitine** | C_10_H_19_NO_4_ | 218.1387 | | 3.79 | | 3 | |
